# Global phylogeography of marine *Synechococcus* in coastal areas reveals strikingly different communities than in the open ocean

**DOI:** 10.1101/2022.03.07.483242

**Authors:** Hugo Doré, Jade Leconte, Ulysse Guyet, Solène Breton, Gregory K. Farrant, David Demory, Morgane Ratin, Mark Hoebeke, Erwan Corre, Frances D. Pitt, Martin Ostrowski, David J. Scanlan, Frédéric Partensky, Christophe Six, Laurence Garczarek

## Abstract

Marine *Synechococcus* comprise a numerically and ecologically prominent phytoplankton group, playing a major role in both carbon cycling and trophic networks in all oceanic regions except in the polar oceans. Despite their high abundance in coastal areas, our knowledge of *Synechococcus* communities in these environments is based on only a few local studies. Here, we use the global metagenome dataset of the Ocean Sampling Day (June 21^st^ 2014) to get a snapshot of the taxonomic composition of coastal *Synechococcus* communities worldwide, by recruitment on a reference database of 141 picocyanobacterial genomes, representative of the whole *Prochlorococcus, Synechococcus* and *Cyanobium* diversity. This allowed us to unravel drastic community shifts over small to medium scale gradients of environmental factors, in particular along European coasts. The combined analysis of the phylogeography of natural populations and the thermophysiological characterization of eight strains, representative of the four major *Synechococcus* lineages (clades I to IV), also brought novel insights about the differential niche partitioning of clades I and IV, which most often co-dominate the *Synechococcus* community in cold and temperate coastal areas. Altogether, this study tackles the main differences between open-ocean and coastal communities worldwide.

## Introduction

Better assessment of the spatial and temporal variability of the genetic diversity, structure and dynamics of marine phytoplankton communities is critical to predicting their future evolution in environments whose physico-chemical properties are continuously altered by the ongoing global change. The marine picocyanobacteria *Prochlorococcus* and *Synechococcus*, together accounting for about 25% of ocean net primary production [1], are key members of phytoplankton communities and constitute particularly relevant models to tackle this issue. *Prochlorococcus* distribution is restricted to the 45°S–50°N latitudinal band preferentially thriving in oligotrophic areas, whilst *Synechococcus* is present in all marine environments from the equator to subpolar waters but reaches its highest abundances in nutrient-rich areas [2–8]. The ability of these two genera to colonize a wide range of ecological niches is likely related to their large genetic diversity [9–13]. For *Prochlorococcus*, numerous environmental and laboratory studies have revealed the clear-cut niche partitioning between physiologically and genetically distinct ecotypes, with ‘phototypes’ [14], ‘thermotypes’ [3, 15, 16], and ‘nutritypes’ [12, 17, 18], occupying distinct light, thermal and nutrient (+Fe/-Fe) niches. Besides *Prochlorococcus*, ‘Cluster 5’ *sensu* [19] also encompasses three major *Synechococcus/Cyanobium* lineages, called sub-clusters (SC) 5.1 through 5.3 [10, 20]. Although a number of phylogenetic studies based on individual markers have considered SC 5.2 and *Cyanobium* as being two distinct lineages (see e.g. [21–23]), the delineation is unclear and it was recently proposed, based on comparative genomics, that all members of these lineages should be gathered into a single group (SC 5.2) named ‘*Cyanobium’*, even though the level of genomic diversity within this group is quite large [20, 24, 25]. SC 5.2 gathers freshwater and halotolerant representatives and thus in the marine environment, members of this group are only found in significant abundance in river-influenced coastal waters, such as the Chesapeake Bay [21, 22, 26] or the Pearl River estuary [23, 27], and in low salinity areas such as the Baltic Sea [28]. SC 5.3 was long thought to contain only obligatory marine representatives and was shown to account for a significant fraction of the *Synechococcus* community in some specific marine areas, including the Mediterranean Sea and northwestern Atlantic Ocean [12, 29–31]. However, freshwater members of this group were recently discovered in the Tous reservoir (Spain) and were then found to be broadly distributed in temperate freshwater lakes [25, 32]. Finally, SC 5.1, a lineage that rapidly diversified after the advent of the *Prochlorococcus* radiation [33, 34], is by far the most widespread and abundant *Synechococcus* lineage in the open ocean environment, e.g. representing more than 93% of total *Tara* Oceans metagenomic reads assigned to SC 5.1-5.3 [12]. From 10 to 15 phylogenetic clades have been defined within SC 5.1 depending on the phylogenetic marker [11, 29, 35] but studies of the global distribution patterns of *Synechococcus* populations in open ocean waters have shown that there are five major clades *in situ* (I, II, III, IV and CRD1), with clades I and IV co-dominating *Synechococcus* communities in cold and temperate, nutrient-rich areas, while clades II, III and CRD1 preferentially thrive in warm waters [6, 12, 30, 31, 36]. Physiological measurement of temperature *preferenda* of strains belonging to clades I, II, III, IV and V isolated across different latitudes further confirmed the existence of warm (clades II, III, V) and cold (clades I and IV) ‘thermotypes’ [37–40]. Despite being phylogenetically distant, clades I and IV were further demonstrated to share a number of physiological adaptations to cold water, including a higher thermal sensitivity of phycobiliproteins [41], a similar change in membrane lipids [40, 42] and an increase of the photoprotection capacities using the orange carotenoid protein (OCP; [43]). Nutrients were also found to play a key role in structuring these populations, with clade II, the most abundant *Synechococcus* lineage in the ocean, dominating the *Synechococcus* community in N-poor areas, clade III in P-poor areas, while CRD1 is restricted to Fe-depleted waters [6, 12, 31, 36].

Although the variability of picocyanobacterial communities and the main physico-chemical factors driving their composition are starting to be well understood in open ocean environments, the picture is much more fragmentary in coastal areas, because only a few coastal sites have been studied to date [21, 22, 24, 27, 44–46]. To get a more global view of the genetic diversity and biogeography of coastal populations of picocyanobacteria, we used metagenomic data from the Ocean Sampling Day (OSD) 2014 campaign [47], encompassing 157 coastal samples collected all over the world at the summer solstice, employing the same protocol for collecting DNA samples and associated metadata. Using a whole genome recruitment (WGR) approach, we assessed the genetic diversity and the clade-level composition of *Synechococcus* communities in OSD samples. Given the previously recognized role of temperature in structuring *Synechococcus* communities, we then analyzed the distribution patterns of the different lineages in light of previously published and new comparative thermophysiological data on *Synechococcus* strains representative of the most abundant clades in the field. The excellent spatial resolution achieved in northern Atlantic and Mediterranean coastal waters allowed us to observe several spatial community shifts and to enlighten the roles of temperature and salinity as key drivers of coastal *Synechococcus* community composition.

## Materials and methods

### Ocean Sampling Day metagenomics data

OSD 2014 is a global sampling campaign that took place on June 21^st^, 2014 and sampled 157 stations worldwide for metagenomes (Dataset S1). The median distance to the nearest coast was 0.29 nautical miles (average: 6.3 nautical miles). Details about sampling methods can be found at https://repository.oceanbestpractices.org/handle/11329/616 [48]. Metagenomic data are available from the European Nucleotide Archive (http://www.ebi.ac.uk/ena/data/view/PRJEB8682) under the study accession number PRJEB8682 (raw data) and from the European Bioinformatics Institute (EBI) Metagenomics portal under the project accession number ERP009703 (processed data). Data were downloaded from the EBI for 150 of the 157 stations for which a “processed reads without annotation” file was available, generated following the EBI analysis pipeline v2.0, available at https://www.ebi.ac.uk/metagenomics/pipelines/2.0. Briefly, Illumina MiSeq paired reads were merged using SeqPrep (https://github.com/jstjohn/seqprep) and trimmed for low quality ends, then sequences with more than 10% undetermined nucleotides were removed using Trimmomatic [49] before discarding reads shorter than 100 nucleotides. Contextual data collected at all OSD stations were retrieved from PANGAEA (https://doi.pangaea.de/10.1594/PANGAEA.854419; [50]) and those used in this study are listed in Dataset S1: only water temperature and salinity data were available in a sufficient number of stations to be used. A map of OSD stations used in this study is available as Fig. S1.

### Taxonomic assignment of metagenomic reads

BLASTN v2.2.28+ [51, 52] was used to align metagenomic reads against a database of 863 complete genomes of aquatic bacteria (Dataset S2), gathering 141 genomes of marine picocyanobacteria and 722 outgroup genomes, the latter including 185 cyanobacterial genomes other than *Prochlorococcus* and marine *Synechococcus* listed in Cyanobase (http://genome.microbedb.jp/cyanobase/) as well as 537 genomes of other aquatic microbes downloaded from the proGenomes database (http://progenomes.embl.de/representatives.cgi). Only best-hit matches (option -max_target_seqs 1) with an e-value below 10^−3^ (-evalue 0.001) were kept, and reads matching outgroup genomes were discarded. Based on BLASTN results, reads aligning over more than 90% of their length on a picocyanobacterial genome were extracted from initial read files, and a second BLASTN was run against a database containing only marine picocyanobacterial genomes with default parameters except for a lower limit on percentage of identity of 30% (-perc_identity 30), a filter on e-value of 10^−2^ (-evalue 0.01) and by selecting the blastn algorithm (-task blastn). BLASTN results were then parsed using the Lowest Common Ancestor method [53]. For each read, BLAST matches with over 80% ID aligned over more than 90% of their length against a reference genome were kept if their BLAST score was within 5% of the best score. Then, the read was attributed to the lowest common ancestor of these matches (i.e., strain, clade, subcluster or genus). Counts of reads assigned to the strain or clade levels were ultimately aggregated by clade. Two additional categories were made for reads that could only be assigned to the level of *Synechococcus* subcluster 5.1 (SC 5.1 in Figures 1 and 3) or even *Synechococcus* genus (*Syn* in Figs 1 and 3).

**Figure 1:**
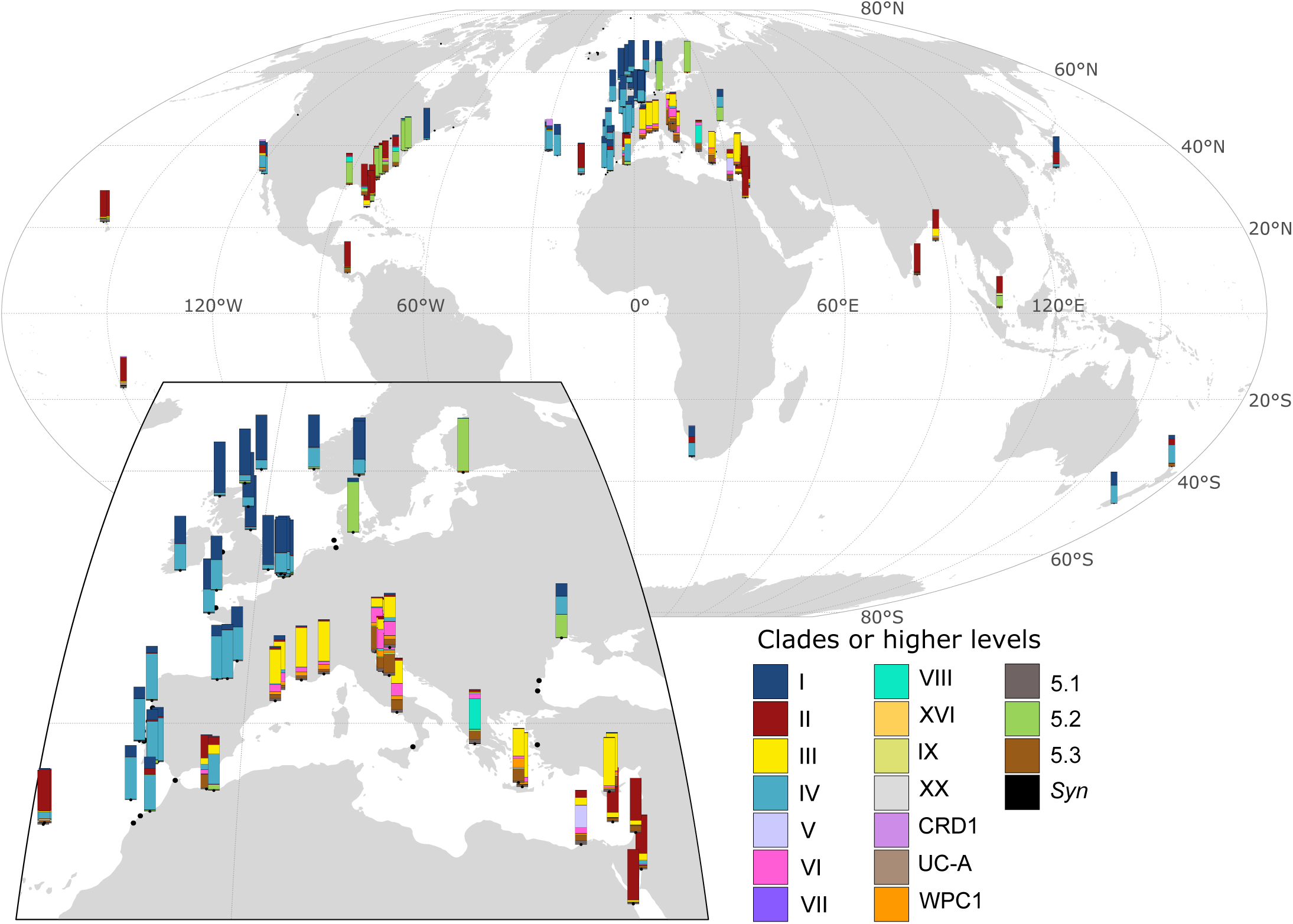
Relative abundance of marine *Synechococcus* clades in OSD stations. Stations are located at the bottom of barplots of relative abundance. The insert shows a close-up version of Europe. Station numbers are shown in Supplementary Figure S1. Categories 5.1 and *Syn* correspond to reads that could not be assigned to a clade but were assigned to the level of *Synechococcus* SC 5.1 or *Synechococcus* genus, respectively.

### Analysis of picocyanobacterial community composition

In order to account for the potential variation in genome length among clades, read counts were divided by the average genome length within each clade. To minimize the noise in recruitment data, we then removed from the dataset stations with less than 600 recruited reads per million bp, corresponding to a genome coverage of ca. 16%, since reads are 242 bp long on average. Read counts at each station were further normalized by the total number of reads recruited at this station to assess relative abundances of taxa. The R packages *cluster* v1.14.4 [54] and *vegan* v2.2-1 [55] were used to cluster stations according to the Bray-Curtis distance. Figures were drawn in R v3.03 with package *ggplot2* v1.0.1 [56].

### Thermal *preferenda* of strains representative of the most abundant clades *in situ*

Two strains of each of the four most abundant *Synechococcus* clades in Fe-replete areas (clades I to IV) were selected from the Roscoff Culture Collection (Table 1; http://roscoff-culture-collection.org/; [57]). Strains were grown in polystyrene flasks in PCR-S11 medium [58] supplemented with 1mM sodium nitrate. The seawater was reconstituted using Red Sea Salts (Houston, TX, USA) and distilled water. Cultures of the eight strains were acclimated at least two weeks to a range of temperatures from 10°C to 33°C, within temperature-controlled chambers (Liebherr-Hausgeräte, Lienz, Austria) and continuous light was provided by green/white/blue LEDs (Alpheus, France) at an irradiance of 20 µmol photons m^-2^ s^-1^. After acclimation, cultures were split into three biological replicates for each strain, and sampled once or twice a day until the stationary phase was reached.

For cell density measurements, aliquots of cultures were preserved with 0.25% glutaraldehyde grade II (Sigma Aldrich, St Louis, MO, USA) and stored at -80°C until analysis [59]. Cell concentration was determined using a flow cytometer (FACSCanto II, Becton Dickinson, San Jose, CA, USA) with laser emission set at 488 nm, and using distilled water as sheath fluid.

To estimate the maximum population growth rates, we considered that *Synechococcus* exponential growth followed:

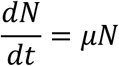

Where *N* is the cell abundances (in cell mL^-1^) and *μ* is the maximum population growth rate (in days^-1^). We estimated *μ* as the coefficient of the linear regression model performed on log-transform *N(t)* data during the exponential phase only.

To overcome the fact that discrete experimental measurements have a limited resolution, we estimated the cardinal growth parameters for each strain using the Cardinal Temperature Model with Inflection (BR model; [60]). This model helps describe the growth response of acclimated phytoplankton strains to temperature using four parameters (Table 2): the optimal temperature for growth (T_opt_) at which the optimal growth rate (μ_opt_) occurs, and the minimal and maximal temperatures for growth (T_min_ and T_max_) at which *μ* = 0. Our data did not allow to constrain T_min,_ but this does not affect our estimation of other parameters.

## Results and Discussion

### Biogeography of coastal picocyanobacterial communities is influenced by seawater temperature

Most of the stations sampled during the OSD 2014 campaign [47] correspond to coastal areas with only 17 of 157 stations located over 11 nautical miles from the nearest coast. This dataset displays a particularly good spatial resolution in some regions of the world ocean and notably along European and Eastern United States coasts, while only a few of the sampled sites were located in the southern hemisphere (7 out of 157; Fig. S1). Here, we used the 150 metagenomes obtained in the framework of this campaign, altogether totaling 41 Gbp (168.7 million reads), to assess the relative abundance of *Synechococcus/Cyanobium* and *Prochlorococcus* clades using a Whole Genome Recruitment (WGR) approach against a reference genome database encompassing 141 genomes of marine picocyanobacteria as well as 722 cyanobacterial or other aquatic microbial genomes, used as outgroups (Fig. 1). *Prochlorococcus* was only abundant at a few stations, likely due to the coastal localization of the sampling sites, and was therefore not included in subsequent analyses. By contrast, *Synechococcus/Cyanobium*, known to outnumber *Prochlorococcus* in coastal areas [2, 8, 24, 61], was detected with sufficient coverage to perform reliable taxonomic assignment at the clade level in 102 out of the 150 OSD metagenomes. At most stations, the *Synechococcus/Cyanobium* community was dominated by one or two taxa among SC 5.1 clades I-IV, SC 5.2 or SC 5.3 (Fig. 1). Consistent with previous studies on the picocyanobacterial distribution in open ocean waters [7, 12, 15, 29, 31, 36], clades I and IV dominated at latitudes above 35°N (except in the Mediterranean Sea) and clade II at latitudes below 35°N, while clade III was almost exclusively present and often dominant in the Mediterranean Sea. It is also worth noting that the co-occurrence of clades I and IV at the few stations beyond 35°S in the Southern hemisphere mirrored the profiles obtained at the same latitude in the Northern hemisphere, in agreement with previous observations in open ocean waters [12, 29, 31, 36] as well as with the low temperatures of isolation sites of clade I and IV strains [38].

In order to further explore the role of temperature on the differential latitudinal distribution of members of clades I to IV, we characterized the thermal *preferenda* of eight strains belonging to these clades (Fig 2). While several strains belonging to clade I were previously shown to withstand colder temperatures than their tropical clade II counterparts [38, 40, 43], growth optima and boundary limits for temperature were only available for one clade IV [40, 62] and two closely related clade III strains [37, 40, 63] and results were obtained in different light conditions, making them difficult to compare. Here, the direct comparison of clades I and IV strains grown in the same conditions showed quite similar thermal preferences. All tested strains displayed an optimal temperature for growth of about 24°C according to our model fit (Fig. 2 and Table 2) and were all able to grow at the lowest tested temperature, 10°C, which is also the lowest temperature measured in the OSD 2014 stations where the *Synechococcus* community was analyzed. In comparison, clades II and III strains were not able to grow at temperatures of 13°C and below, thus confirming with several strains that clades I and IV are cold thermotypes, whereas clades II and III are warm thermotypes. Altogether, these results support the idea that differences in thermophysiology at least partially explain the latitudinal distribution of these four clades.

**Figure 2:**
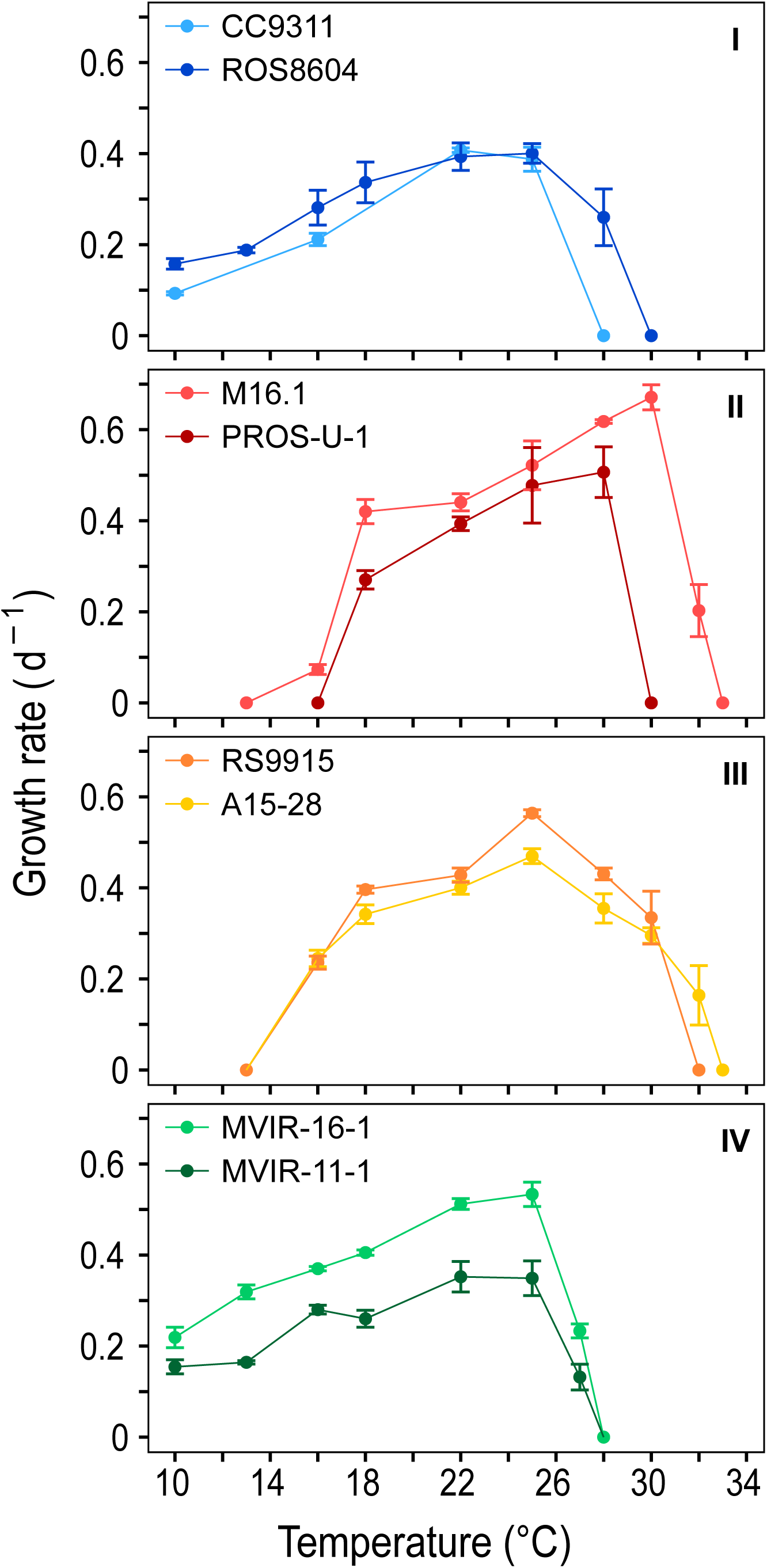
Temperature preferenda of eight marine *Synechococcus* strains. Growth rate as a function of temperature of acclimated growth. Two strains were chosen within each of the four major clades I, II, III and IV (top to bottom). All cultures were grown at a light intensity of 20 µmol quanta m^-2^ s^-1^. Error bars are standard deviation from the mean based on at least 3 replicates (n≥3).

Besides the abundance of clades I and IV, coastal *Synechococcus* communities also exhibited some other specificities as compared to open ocean populations, notably the very low relative abundance of clade CRD1, which was shown to be prevalent in large regions of the open ocean that are limited by iron availability [12, 31, 36], as well as the dominance of SC 5.2 in the brackish Baltic sea and at stations along the Atlantic coast of North America, often co-occurring with a low proportion of clade VIII. The latter observation is most likely due to the influence of riverine inputs at these OSD stations, these taxa being known to occur in estuarine areas and to contain strains growing over a large range of salinity [10, 21, 35]. This hypothesis was further confirmed by clustering stations according to the relative abundance profiles of *Synechococcus* clades (Fig. 3), which clearly separated stations dominated by subcluster 5.2 and showed that they had a lower salinity than most other stations (cluster 5, Fig. 4B). Finally, clades V and VI, which were not distinguished from clade VII (and CRD1) in previous global surveys of *Synechococcus* distribution using the 16S rRNA marker gene, were found to be locally abundant in the dataset. While the V/VI/VII/CRD1 group was considered to be widely distributed in oceanic waters [4, 15, 29], our analysis reveals the potential preference for coastal areas of the closely related clades V and VI. This result is consistent with the previous local observations of the occurrence of clade V- and VI-related sequences at some coastal sites in the Adriatic Sea and the Pearl River Estuary [23, 64, 65].

**Figure 3:**
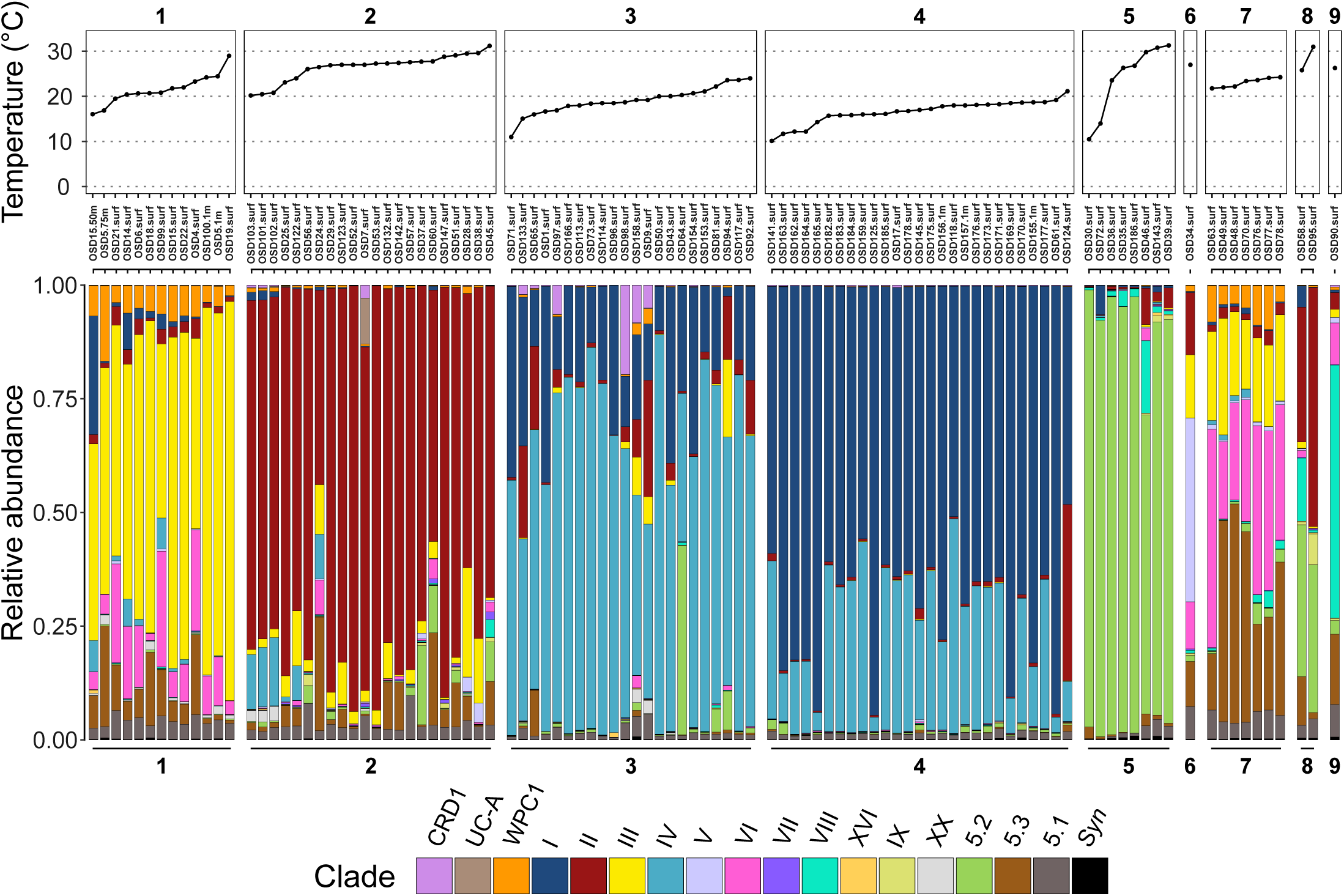
Clusters of OSD stations based on relative abundance profiles of *Synechococcus* clades. OSD stations were clustered based on the relative abundance profiles of marine *Synechococcus* clades using Bray-Curtis distance. The upper panel indicates water temperature. Categories 5.1 and *Syn* correspond to reads that could not be assigned to a clade but were assigned to the level of *Synechococcus* SC 5.1 or *Synechococcus* genus, respectively.

**Figure 4:**
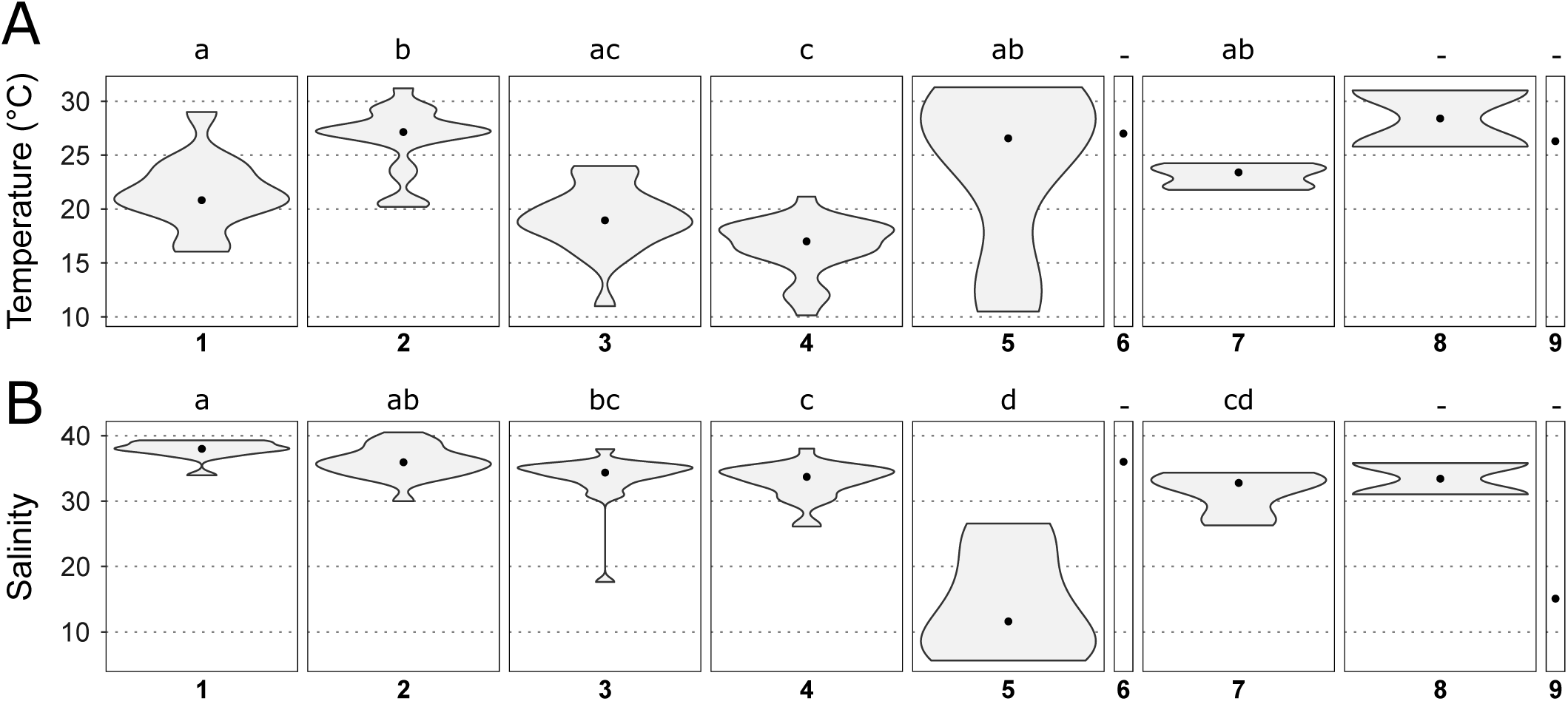
Violin plots showing the distribution of temperature and salinity for each cluster of OSD stations defined in Fig. 3. A. Temperature. B. Salinity. The black dot in each violin plot shows the median value. Different letters indicate significantly different distributions (Dunn test. adjusted p-value < 0.05). The same analysis considering distance to the nearest coast gave no significant result.

### A progressive latitudinal shift in *Synechococcus/Cyanobium* communities along the coast of Europe

Besides the abovementioned specificities of coastal regions in terms of *Synechococcus/Cyanobium* community composition, we also observed changes in communities at a finer spatial scale along European coasts, where the sampling effort was the highest (see zoom in Fig. 1 and Fig. S1 for station numbers). While along the southern part of this latitudinal gradient from the Moroccan to French Atlantic coasts, *Synechococcus* communities were dominated by clade IV (e.g. OSD92), a progressive northward shift was observed towards the dominance of clade I in the North Sea (e.g., OSD164). Clustering of stations based on clade relative abundance indeed highlighted two groups of stations, the first one dominated by clade IV (cluster 3) and the second one by clade I (cluster 4; Fig. 3). Interestingly, clade I was found to dominate at stations that display a significantly lower salinity than those dominated by clades II or III (clusters 1 and 2). These clade I-dominated stations also exhibited a significantly lower temperature (average 16.6°C, median 17°C) than all other clusters except cluster 3 dominated by clade IV (average temperature 19.1°C, median 19°C), the latter cluster of stations showing a significant difference in temperature only with cluster 2 (dominated by clade II). Thus, despite a clear latitudinal shift in the ratio of clade I to clade IV along the European coast, neither the difference in salinity nor the difference in seawater temperature seem to be sufficient to fully explain the observed changes.

While our thermal *preferenda* measurements confirmed that clades I and IV are both cold thermotypes, strains used for this study showed similar growth rates at cold temperatures (Fig 2). This supports the observation that temperature alone does not explain their differential distribution, with the caveat that only two strains per clade were analyzed. As previously suggested for clade I in a number of environmental studies [5, 7, 23, 66, 67], it is possible that clade I and potentially clade IV are comprised of distinct genotypes exhibiting different lower temperature boundary limits and colonizing different thermal niches. A previous study indeed showed variability in the minimal growth temperature of clade I strains in relation to their latitude of isolation [38], and comparison of our experimental data with previous data acquired under the same light conditions [40] brings evidence of such variability for clade IV. Indeed, the two clade IV strains characterized here were sampled at high latitude (Table 1) and show a higher tolerance to cold temperatures than BL107, another clade IV strain isolated in the Mediterranean Sea [40]. Thus, the ecological differences between clades I and IV are most probably difficult to identify due to underlying differences between genotypes at a finer taxonomic level, and a higher taxonomic resolution would be necessary if one wanted to observe a significant effect of temperature on the distribution of populations of these clades. Alternatively, it is possible that other parameters or combinations of parameters varying with latitude need to be considered to explain the shift in clades I and IV dominance. Notably, an interaction between light and temperature on phytoplankton physiology has been described in a number of species [68], and these two parameters vary greatly with latitude.

Several other potential reasons have been previously evoked to explain the co-occurrence of clades I and IV and variations in their relative abundance *in situ*, including differences in their metal concentration requirements [31, 44] and transport and mixing of populations by advection. The latter hypothesis was notably suggested for *Synechococcus* populations from north-west of the Svalbard island (above 79°N), where the Gulf Stream current was proposed to be the source of clade IV populations in summer [5] and from the Korean Sea where the warm, oligotrophic Kuroshio Current was suggested to be responsible for the co-occurrence of clades I, II and IV populations [69]. Interestingly the only OSD sampling site close to Japan (OSD124, Fig. S1) was described by its sampler as a site where oceanic and coastal waters are sporadically interchanged, which could explain the unexpected profile of this station where clades I and II co-dominate (Fig. 1). Finally, other studies also suggested that clade I could be a more coastal and opportunistic clade than clade IV [10, 44], but this hypothesis does not seem to be confirmed by the present study since many coastal stations (cluster 3) are actually dominated by clade IV.

### Local changes in *Synechococcus* communities in the Mediterranean Sea

Stations sampled in the Mediterranean Sea fell into several clusters based on their composition in *Synechococcus/Cyanobium* lineages. Most stations belonged to cluster 1, dominated by clade III with a low relative abundance of clades VI, WPC1 and SC 5.3 (Fig. 3). This clade composition is quite similar to that previously described [12] for open waters of the Mediterranean Sea, which was suggested to be related to specific features of this semi-enclosed sea and notably to its low phosphate concentration [12, 15, 29], a parameter that was not available in the OSD dataset. Most of the stations of the Adriatic Sea formed a distinct cluster (cluster 7), where the same clades were present but in different proportions, clade VI and SC 5.3 taking over clade III. Finally, stations OSD34 and OSD90, located on the Egyptian and Greek coasts, respectively, the only stations of the OSD dataset comprising a high proportion of clade V or VIII, formed a cluster on their own (cluster 6 and 9). While these four clusters (clusters 1, 6, 7 and 9) are specific to the Mediterranean Sea, it is worth noting that two stations at the easternmost end of the Mediterranean Sea (OSD123 and OSD132, Figs. 3 and S1) fell into cluster 2, dominated by clade II, and showed a clade composition very similar to the samples collected in the Red Sea (OSD52 and OSD53). This suggests that Israeli coastal areas are strongly influenced by waters entering the Mediterranean Sea via the Suez Canal, consistent with previous findings for *Synechococcus, Prochlorococcus* as well as for larger organisms [70, 71].

Interestingly, the three specific clusters identified in the Mediterranean Sea displayed different temperature and salinity characteristics (Fig. 4A-B). The salinity range of stations in cluster 1 (dominated by clade III) was narrow (average salinity 37.90 psu, median 37.98 psu) and significantly higher than that of cluster 7 (dominated by clade VI and SC 5.3, average salinity 31.43 psu, median 32.77 psu), suggesting that clade VI and SC 5.3 are able to cope with lower salinities. Consistently, SC 5.3 was recently found to encompass members colonizing freshwater lakes [25, 32], while in the marine environment, this subcluster was reported both in strictly marine waters [12, 30] and in low salinity waters [72]. Our study also brings new insights into the ecological niche occupied by clade VI, whose distribution was so far poorly known [29], and that appears to be restricted to coastal regions of intermediate salinity. All stations of the Adriatic Sea comprising cluster 6 were indeed sampled in the northwestern part of this area, where the influence of the Po River plume may be important [73]. This distribution is consistent with previous observations of the closely-related and often co-occurring clade V in low salinity surface waters of the Adriatic Sea [64] and of both clades V and VI in the Pearl River Estuary [74]. Laboratory experiments also showed that representative strains of these two clades can tolerate salinities as low as 15 psu [75]. Still, we cannot exclude that besides low salinity, other local specificities linked to riverine input might also explain the predominance of SC 5.3 and clade VI in coastal areas of the Adriatic Sea.

A significant difference in water temperature was also found between cluster 1, dominated by clade III (average temperature 21.5°C, median 20.8°C) and cluster 2, dominated by clade II (average 26.5°C, median 27.1°C). This suggests that the shift observed at the easternmost part of the Mediterranean Sea from a dominance of clade III to a local dominance of clade II (stations OSD123 and OSD132, Figs. 1 and S1) might be related to a difference in water temperature. Interestingly, in contrast to clades I and IV that often co-occur, clades II and III seem to be nearly mutually exclusive, at least in the Mediterranean Sea, and the temperature limit above which clade II dominates seems to lie around 25°C (Fig. 3). In our experimental comparison of thermal *preferenda*, this corresponds to the temperature at which growth rates of clade II strains become higher than that of clade III strains, resulting in a higher optimal temperature of clade II compared to clade III strains (Table 2). Altogether, temperature and salinity appear as major factors driving the composition of *Synechococcus/Cyanobium* communities in coastal waters of the Mediterranean Sea, although other biotic and abiotic factors are most likely involved, notably the availability of phosphorus, a key limiting nutrient in this area [76].

## Conclusion

The OSD dataset is unique, not only by providing an instantaneous snapshot of the microbial community composition but also because, by focusing on coastal areas, it nicely complements other recent global ocean surveys performed in the open ocean [6, 12, 31, 36, 77, 78]. In particular, the good spatial resolution of the sampling performed along the European coasts is well-adapted to observe shifts in communities and delineate their boundaries. Despite the fact that only a few physico-chemical parameters were collected, this dataset allowed us to considerably improve our knowledge of the distribution of *Synechococcus/Cyanobium* lineages in coastal areas, to gain insights into the realized environmental niches of the main ones, including some that were previously poorly known such as clade VI, as well as to reinforce hypotheses about thermal niche differentiation that were supported by laboratory experiments on a set of representative strains. A continued effort towards global instantaneous surveys of microbial diversity in coastal areas over the long term and at different seasons would be invaluable to monitor the evolution of microbial communities in relation to global change.

## Acknowledgements

We thank the OSD Consortium for sampling, sequencing and making freely available the data analyzed in this paper as well as the Roscoff Culture Collection (http://roscoff-culture-collection.org/) for providing *Synechococcus* strains used in this study. Financial support for the OSD program was provided by the European Union program MicroB3 (UE-contract-287589) and authors were supported by the French “Agence Nationale de la Recherche” programs SAMOSA (ANR-13-ADAP-0010) and CINNAMON (ANR-17-CE02-0014-01) as well as the European program Assemble Plus (H2020-INFRAIA-1-2016-2017; grant no. 730984). DJS received funding from the European Research Council (ERC) under the European Union’s Horizon 2020 research and innovation programme (grant agreement No 883551).

## Supplementary information

**Supplementary Figure S1**: **Map of OSD stations**. All OSD stations that were analyzed in this study are indicated by their number. The inset shows a close-up view of Europe.

**Dataset S1: OSD samples used in this study**. Characteristics and accession numbers of the OSD samples analyzed in this study and corresponding contextual data, as retrieved from PANGAEA (https://doi.pangaea.de/10.1594/PANGAEA.854419; [50]).

**Dataset S2**: **Summary data for the 863 complete genomes of aquatic bacteria used as reference in this study**. Genomes sequences were retrieved either from Cyanorak *v2*.*1* (www.sb-roscoff.fr/cyanorak), NCBI Genbank for additional *Synechococcus* whole genomes and for genomes other than marine *Synechococcus* and *Prochlorococcus* listed in Cyanobase (http://genome.microbedb.jp/cyanobase/), or proGenomes (http://progenomes.embl.de/index.cgi). The table includes subclade designation based on [11].

