## Supplemental Figure 1 for "Global phylogeography of marine *Synechococcus* in coastal areas reveals strikingly different communities than in the open ocean"

### Supplementary Figure

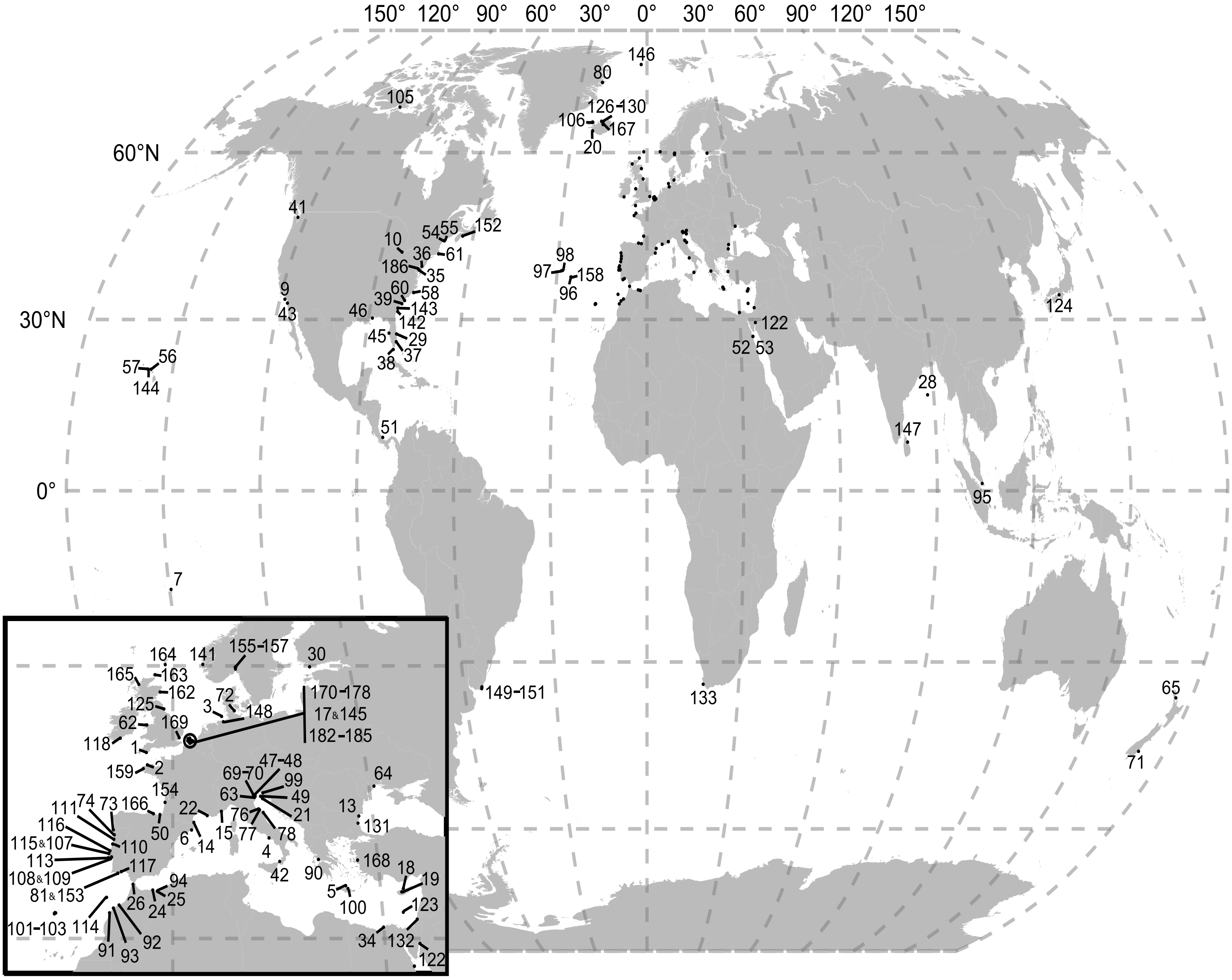


**Supplementary Figure S1**: **Map of OSD stations.**
All OSD stations that were analyzed in this study are indicated by their number.
The inset shows a close-up view of Europe.
